# Validating Dynamicity in Resting State fMRI with Activation-Informed Temporal Segmentation

**DOI:** 10.1101/2020.10.12.335976

**Authors:** Marlena Duda, Danai Koutra, Chandra Sripada

## Abstract

Confirming the presence (or absence) of dynamic functional connectivity (dFC) states during rest is an important open question in the field of cognitive neuroscience. The prevailing dFC framework aims to identify dynamics directly from connectivity estimates with a sliding window approach, however this method suffers from several drawbacks including sensitivity to window size and poor test-retest reliability. We hypothesize that time-varying changes in functional *connectivity* are mirrored by significant temporal changes in functional *activation,* and that this coupling can be leveraged to study dFC without the need for a predefined sliding window. Here we introduce a straightforward data-driven dFC framework, which involves informed segmentation of fMRI time series at candidate FC state transition points estimated from changes in whole-brain functional activation, rather than a fixed-length sliding window. We show our approach reliably identifies true cognitive state change points when applied on block-design working memory task data and outperforms the standard sliding window approach in both accuracy and computational efficiency in this context. When applied to data from four resting state fMRI scanning sessions, our method consistently recovers five reliable FC states, and subject-specific features derived from these states show significant correlation with behavioral phenotypes of interest (cognitive ability, personality). Overall, these results suggest abrupt whole-brain changes in activation can be used as a marker for changes in connectivity states, and provides strong evidence for the existence of time-varying FC in rest.

## 1. Introduction

Over the past two decades the study of functional connectivity (FC) has emerged as a preeminent method in cognitive and clinical neuroscience, aiming to characterize the functional network organization of the brain, and to identify objective markers of neuropsychiatric diseases and clinically relevant phenotypes. FC describes the temporal correlation in activation patterns of spatially distinct regions of the brain, typically measured by blood oxygen level-dependent (BOLD) functional magnetic resonance imaging (fMRI). Originally, the entire field of FC was built on a critical assumption: that patterns of connectivity are static during any given measurement interval in a resting state, i.e. the absence of any cognitive task (Biswal et al., 1995). Static FC has been used to identify global differences in functional network organization of the brain between cognitive task states and resting state (Greicius et al., 2003), as well as to characterize differences in FC between healthy controls and subjects with neuro-psychiatric diagnoses, such as schizophrenia (Lynall, 2010) or autism spectrum disorder (Hull et al., 2017).

Recently, however, a number of studies have questioned this assumption, instead advocating the “dynamic” or “time-varying” connectivity view that functional connectivity patterns exhibit substantial moment-to-moment changes over time, specifically within a standard fMRI measurement interval of five to fifteen minutes (Calhoun et al., 2014; Chang & Glover, 2010; Cohen, 2018; Hutchison et al., 2013; Lurie et al., 2019; Preti et al., 2017). These changing FC patterns are thought to correspond to cognitively meaningful discrete FC network configurations, or connectivity states, that are reproducible both within and between individual subjects. Dynamic states have been documented across different populations including children (Marusak et al., 2018) and adults (Allen et al., 2014; Cai et al., 2018; Chen et al., 2016; Choe et al., 2017; Liu & Duyn, 2013; Smith et al., 2018), and have been supported with concurrent electroencephalography (EEG) data (Allen et al., 2018; Chang et al., 2013; Tagliazucchi et al., 2012). Furthermore, it has been shown that other characteristics such as the amount of time spent in specific states and the number of transitions between states vary with meaningful individual differences such as age (Cabral et al., 2017; Hutchison & Morton, 2015; Marusak et al., 2016), sex (Mao et al., 2017), or disease status (Cordes et al., 2018; Damaraju et al., 2014; Jones et al., 2012; Rashid et al., 2014).

By definition, the presence of dynamic functional connectivity (dFC) in resting state is marked by changes in the connectivity structure of the fMRI time series. The prevailing sliding window framework aims to identify these second-order changes using functional connectivity “snapshots” obtained from time windows of fixed length slid across the entire fMRI time series. The resultant windowed connectomes are then flattened into feature vectors, concatenated across subjects, and clustered into *k* distinct connectivity states. The sliding window approach represents an important advance in the study of time-varying brain connectivity, but it nonetheless suffers from several important limitations.

First, the sliding window method relies heavily on the arbitrary choice of window size, and results can differ substantially across various window widths (Hindriks et al., 2016; Shakil et al., 2016). A second problem is that simulations suggest that sliding window methods can introduce artifactual connectivity variation even under conditions when such variation is known to be absent (Laumann et al., 2017; Lindquist et al., 2014). Third, perhaps due to one or more of the preceding issues, the sliding window method has been found to have poor test-retest reliability (Choe et al., 2017). Fourth, the overlapping nature of the sliding windows precludes definitive segmentation of the fMRI time series into states, making interpretation of the state dynamics difficult. Finally, the sliding window approach requires constructing a sizable number of overlapping windowed connectivity matrices: with 400 timepoints and a 30 second window, 370 distinct connectivity matrices are required (at a TR = 1s). This poses serious scalability issues for relatively long or more temporally granular fMRI datasets.

Some alternatives to sliding window approaches have been proposed in recent years; however, these too have certain drawbacks and limitations. The dynamic conditional correlation (DCC) model is a multivariate volatility model that estimates the changing covariance structure at each timepoint in the fMRI time series (Choe et al., 2017; Lindquist et al., 2014). While the DCC model allows for a parametric approach to estimating framewise FC with robust statistical inference, it increases the number of connectivity matrices to consider in the final clustering step compared to the sliding window method, further hindering its scalability. Furthermore, the formulation of the DCC model has been shown to give biased results in high dimensional data (Hafner & Reznikova, 2012), which poses an issue for application in fMRI data with ROIs and time points. Hidden Markov models (HMMs), which seek to decompose a time series into a sequence of discrete “hidden” states, are another increasingly popular approach for estimating connectivity dynamics (Baker et al., 2014; Quinn et al., 2018; Vidaurre et al., 2017; Zhang et al., 2020). However, HMMs rely on several strong assumptions including a predefined number of *k* hidden states that transition between one another in a Markovian fashion (state transitions depend solely on the state at the previous time point). Moreover, HMMs trained at the group level assume a single governing state-to-state transition structure across all subjects, which may be too strict and miss important individual variability.

Our focus here is on a windowless alternative to the sliding window that simplifies the problem of testing for dynamicity in a resting state as much as possible. It is well known from the task-based fMRI literature that task-driven changes in activation patterns co-occur with changes in connectivity patterns (Sripada et al., 2014; Davison et al., 2015; Gonzalez-Castillo et al., 2015; Spielberg et al., 2015; Telesford et al., 2016; Shine & Poldrack, 2018). This coupling of activation and connectivity changes suggests the possibility that changes in the *activation structure* of the fMRI time series, which are easily derived, can serve as a reasonably reliable marker for changes in the *connectivity structure,* which are more difficult to obtain in an unbiased way. Though connectivity changes may not always be accompanied by activation changes, as long as there is significant correspondence, we can leverage the latter (straightforwardly identified) to find the former (less so) without the need to define a window of arbitrary length.

In this work we leverage the coupling between activation and connectivity to present the activation-informed segmentation approach, a straightforward dFC framework centered around informed segmentation of fMRI time series at candidate FC state change points. Moment-to-moment changes in functional activations have previously been utilized in the literature to investigate dynamic functional connectivity (Shine et al., 2015), but have yet to be used to localize connectivity state changepoints for dynamic time series segmentation. Our simple and intuitive approach detects significant instantaneous changes in functional activation patterns and generates data-driven segments of stable connectivity throughout the fMRI time series. For clarity, we will use the term “segments” when referring to our method and “windows” when referring to the sliding window approach. Separating the time series into discrete time segments rather than highly overlapped sliding windows significantly improves the computational efficiency of dFC analysis and enhances interpretability of results by enabling precise identification of state transition junctures—something the sliding window method cannot provide. We suggest that these FC-tailored segments provide a useful alternative to standard sliding windows in dFC analyses and show that our approach significantly outperforms the sliding window paradigm in recovering known FC state transitions in a block-design task. Furthermore, we propose a framework for the comparison of connectomes derived from segments of variable length, as well as a graph embedding step for summarizing connectomes into low-dimensional representations that we show are better suited for downstream clustering and machine learning tasks than current approaches.

## 2. Methods

### 2.1. Data Description

#### 2.1.1. HCP Data

In this work, we utilize the Human Connectome Project (HCP) S1200 Young Adult dataset made publicly available through the Washington University and the University of Minnesota HCP consortium (http://humanconnectome.ora). It is one of the richest collections of neuroimaging data to date, consisting of structural and functional MRI, behavioral assessments and genotypes for 1200 healthy subjects ages 22–35. A full description of the acquisition protocol can be found in (Van Essen et al., 2013). In short, all HCP fMRI data were acquired on a modified Siemens Skyra 3T scanner using multiband gradient-echo EPI (TR = 720 ms, TE = 33 ms, flip angle = 52°, multiband acceleration factor = 8, 2 mm isotropic voxels, FOV = 208 × 180 mm, 72 slices, alternating RL/LR phase encode direction). Participants completed four total resting state fMRI scanning sessions (two sessions collected on each of two different days). Each resultant resting state fMRI time series consisted of 1200 volumes sampled every 0.72 seconds, for a total acquisition time of 14 minutes and 24 seconds.. During the resting state sessions participants were instructed to keep their eyes open and fixated on a cross hair on the screen, while remaining as still as possible. For clarity, we will refer to resting state data from the first collection day as sessions 1A (RL) and 1B (LR), and similarly sessions 2A and 2B for those collected on the second day.

Though our main objective is to assess FC dynamics during rest, we also leverage the repeating task/rest block structure of the working memory (WM) task data available in HCP as a natural ground truth to test the performance of our method in identifying the known transitions between the task and rest conditions. The HCP WM task consists of four repeating task/rest blocks, where each block is structured as follows: 27.5 seconds Task 1 (0-back), 27.5 seconds Task 2 (2-back), 15 seconds rest. Using the same acquisition details outlined above, each WM task fMRI time series consisted of 405 volumes sampled every 0.72 seconds, for a total acquisition time of 4 minutes and 52 seconds. Two sessions of WM task fMRI were acquired back-to-back, alternating between RL and LR phase encoding directions. We will refer to these as WM session 1 (RL) and WM session 2 (LR).

#### 2.1.2. Data Preprocessing

Processed volumetric data from the HCP minimal preprocessing pipeline including ICA-FIX denoising were used. Full details of these steps can be found in (Glasser et al., 2013; Salimi-Khorshidi et al., 2014). Briefly, BOLD fMRI data were gradient-nonlinearity distortion corrected, rigidly realigned to adjust for motion, fieldmap corrected, aligned to the structural images, and then registered to MNI space with the nonlinear warping calculated from the structural images. Then FIX was applied on the data to identify and remove motion and other artifacts in the timeseries. These files were used as a baseline for further processing and analysis (e.g. MNINonLinear/Results/rfMRI_RESTl_RL/rfMRI_RESTl_RL_hp2000_clean.nii.gz from released HCP data).

Images were smoothed with a 6 mm FWHM Gaussian kernel, and then resampled to 3 mm isotropic resolution. This step as well as the use of the volumetric data, rather than the surface data, were done to allow comparability with other large datasets in ongoing and planned analyses that are not amenable to surface-based processing. The smoothed images then went through a number of resting state processing steps, including motion artifact removal steps comparable to the type B (i.e., recommended) stream of (Siegel et al., 2017). Further details on motion artifact removal can be found in (Sripada et al., 2019). Lastly, we calculated spatially-averaged time series for each of the 268 ROIs from the parcellation given in (Finn et al., 2015).

For our analysis, we first considered the set of 966 subjects listed in (Sripada et al., 2019) that met the following criteria: structural T1 and T2 data, four complete resting state fMRI sessions, and < 10% of resting state frames censored due to excessive motion (framewise displacement of 0.5 mm). From this set 922 subjects also had two complete WM task fMRI sessions, defining our final subset of subjects.

### 2.2. The Activation-Informed Segmentation Framework

Here we propose a simple framework for assessing dynamic changes in functional connectivity in fMRI time series, termed the activation-informed segmentation method. This method leverages the coupling between changes in connectivity structure and changes in whole-brain activation patterns to produce an intuitive, interpretable and computationally efficient alternative to the sliding window approach. Our framework consists of three main steps: tailored segmentation of all fMRI time series, summarization of the functional connectivity within each discovered window, and finally segregation and characterization of a final set of connectivity states (Figure 1). These steps are detailed in subsections 2.2.1 - 2.2.3 below.

**Figure 1.**
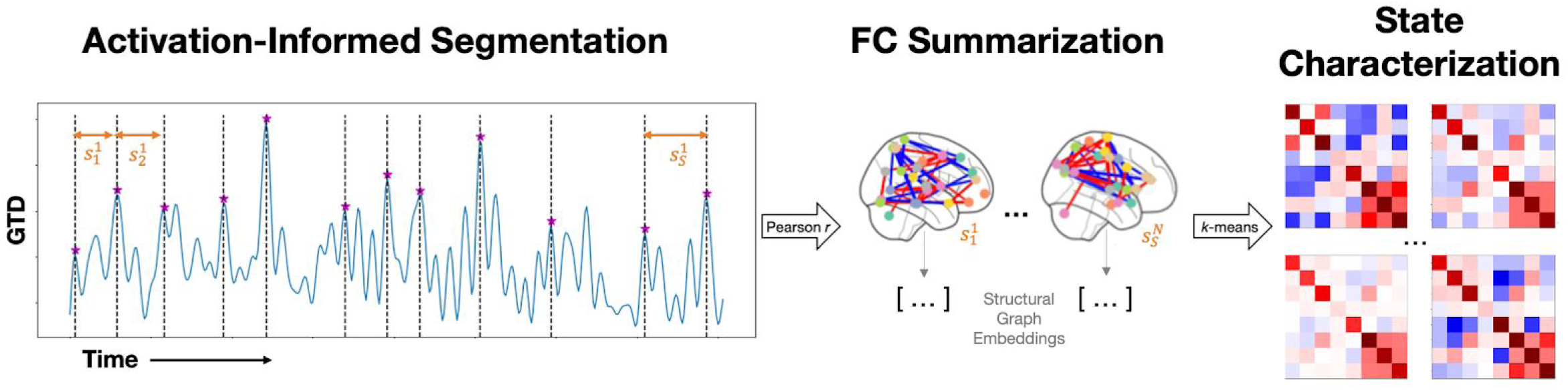
Depiction of our activation-informed segmentation pipeline. Briefly, peaks in the GTD series define the boundaries of our tailored, non-overlapping stable-FC segments *S*_1_ to *S_S_* (note *S* can vary between subjects) for all subjects *I−N*. Next, functional connectivity is summarized using structural graph embeddings for each segment in the set of all segments 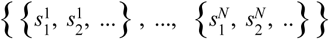. Finally, *k*-means is applied to segregate all segments into a set of *k* connectivity states.

#### 2.2.1. Activation-informed time series segmentation

The dynamic FC paradigm suggests the presence of significant instantaneous changes in connectivity structure at transition points between two distinct functional states. Using this logic, we sought to identify potential connectivity state change points within fMRI data and utilize them to perform informed segmentation of the time series as a metric for assessing FC dynamics. To estimate the changes in functional connectivity from one time point *t* to the next, we observe changes in functional activation from one time point to the next by calculating the temporal derivative *(dt)* of each of *n* ROI time series *(ts)* of length *T* using first-order differencing similar to that in the multiplication of temporal derivatives (MTD) method (Shine et al., 2015):

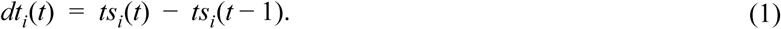

At this point, our method importantly diverges from the MTD method: while the MTD uses these granular temporal derivatives to define the connectivity between each pair of ROIs, our method further summarizes them to provide an estimate of moment-to-moment changes in activation on the whole brain scale. The resulting *n* temporal derivative series of length *T-1* are then summarized by taking the L2-norm at each time step *t.* resulting in a single vector of length *T-1,* which we have termed the Global Temporal Derivative (GTD) series:

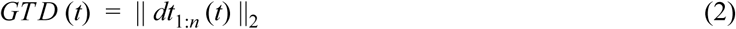

The GTD provides a univariate summarization of instantaneous changes in global brain activation throughout an fMRI time series, therefore peaks in the GTD series correspond to instances of significant moment-to-moment alterations in functional activity. Growing research suggests the global signal is not noise, and carries meaningful information about mental states (Wong et al., 2013). Here, we build on this work to suggest that global signal shifts mark changes in dynamic mental states. We seek to automatically identify these change points as candidate FC state change points for our time series segmentation step. We begin by applying exponentially-weighted moving average smoothing to the GTD series to reduce noisy peaks. We then perform moving average peak detection on the smoothed GTD series, identifying points in the time series that are >=2.5 standard deviations above the moving average. To avoid identification of multiple points that surpass this threshold but actually correspond to a single true peak, we collapsed points in close proximity to one another to the local maximum (within 10 TR, corresponding to 7 seconds or the approximate time-to-peak of the hemodynamic response function (Friston, 2003)). Furthermore, as these change points define our tailored segments for downstream calculation of functional connectivity we set a minimum inter-peak distance of 25 TR to ensure sufficiently large segments for calculating Pearson correlation (Schönbrodt & Perugini, 2013; Thirion et al., 2007; Turner et al., 2018) (note: we reduce this to 15 TR for the case of WM task data to accommodate the shorter resting state segments we intend to capture). This final set of change points define the boundaries of the tailored time segments, within which we compute FC and between which we investigate potential dynamic FC shifts.

#### 2.2.2. Functional Connectivity Estimation

For each tailored segment *s*, we compute the functional connectivity matrix *C*^*(s)*^, where the *i,j*^th^ entry is the Pearson correlation of ROIs *i* and *j* within the time segment, *ts* (*s*):

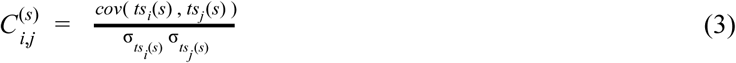

We then apply the Fisher transformation followed by z-scoring on each FC matrix *C*^*(s)*^, to allow for better comparisons between connectivity matrices of segments of differing lengths. Connectivity matrices derived from shorter segments have, on average, higher correlation values than those from longer segments, resulting in a skewed sample distribution. Applying the Fisher transformation enforces an approximately normal distribution of the connectivity values within each segment (Fisher, 1915), and the z-score then translates these connectivity values in terms of their standard deviations from the mean. While these connectome transformations are common practice in the field of FC, they are especially important when attempting to compare connectomes from segments of variable lengths, which is illustrated in Figure 2.

**Figure 2.**
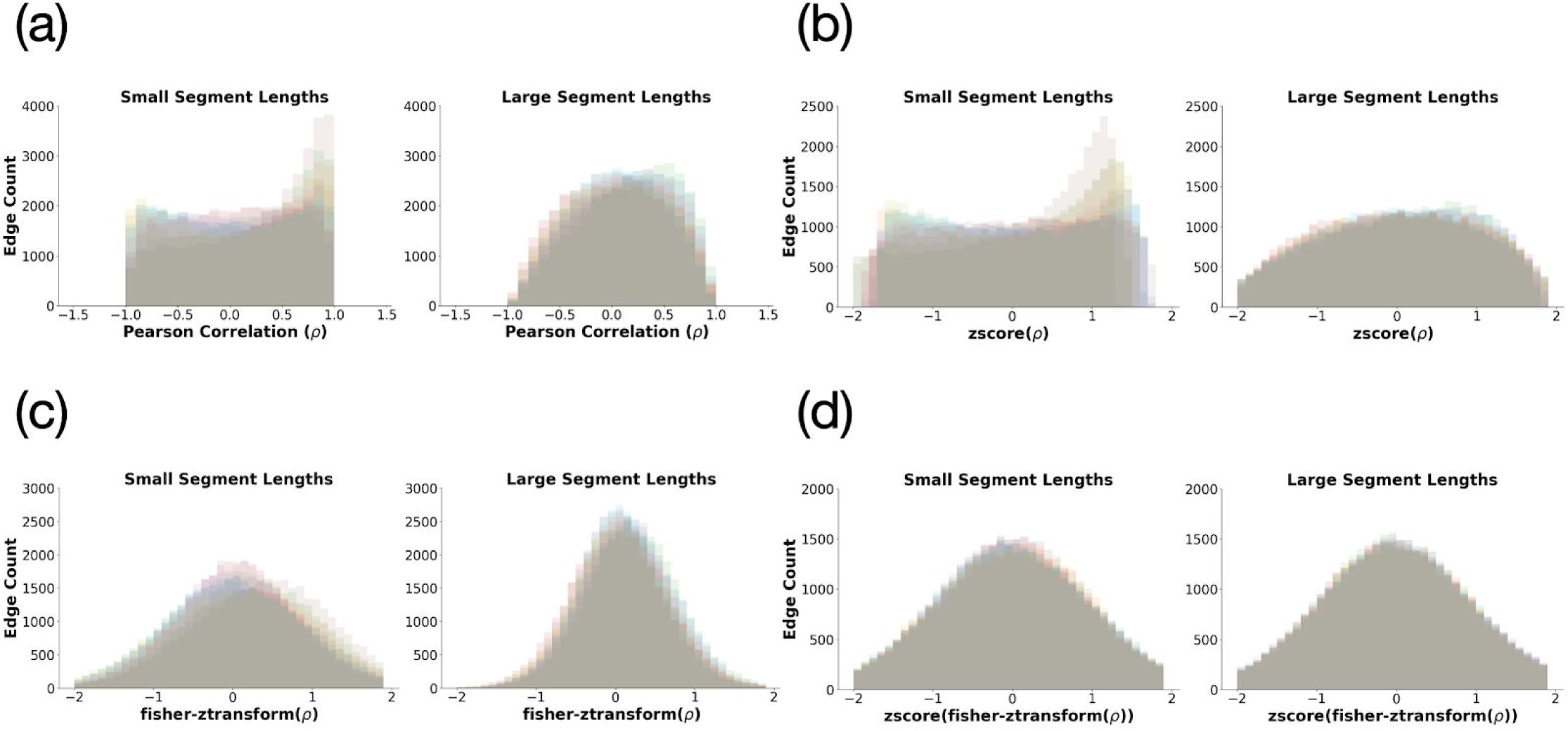
Effects of (A) no transformation, (B) z-score transformation, (C) Fisher transformation and (D) z-scored(Fisher) transformation on the distribution of Pearson correlation-based connectivity values in short (< 25 TR) and long (> 35 TR) segments.

Thresholding is another common pre-processing step in functional connectivity analysis, as it preserves only the high-fidelity connections within connectomes and effectively filters out noise. Though the Fisher transformation with z-scoring helps to align the sample distributions of connectivity values between longer and shorter segments, we still observed the effects of segment length when thresholding on z-scores alone—connectomes from shorter segments were denser (i.e. had more edges preserved) after thresholding than connectomes from longer segments. This segment-length discrepancy in connectome density with z-score thresholding had significant downstream effects in our pipeline, as we found the resultant FC state clusters were highly correlated with segment length. To avoid these segment length effects, we fix the density of all connectomes by thresholding to the top-*K* connections (or edges) in each connectome. Here, we set top-*K* = 10,000, which preserves the strongest (i.e. highest magnitude) 27.95% edges, thereby providing sufficient noise reduction.

#### 2.2.3. State Clustering

The final step of our dFC framework involves using *k*-means clustering to separate all thresholded connectomes into a discrete set of *k* connectivity states. This state clustering occurs on the aggregated set of *m* connectomes, where *m* is the total number of time segments across all subjects in a single fMRI scanning session (Table 1). In traditional dFC streams, this approach involves performing means clustering on the flattened upper triangular of all *m* connectomes, however we found poor performance with this method, likely due to the high dimensionality of the flattened connectomes (>35,000) (Supplemental Table 1). We address this issue of high dimensionality by generating low-dimensional latent representations of each thresholded connectivity matrix that sufficiently summarize the connectivity patterns within the time segment. Specifically, we utilize state-of-the-art graph embedding methods, which are commonly used in the field of data mining to generate low-dimensional representations of graphs (i.e. networks) (Rossi et al., 2020). Connectomes are graphs by definition, consisting of a set of nodes (ROIs) connected by edges (z-scored correlations), so graph mining methods naturally extend to the connectome space. To generate our graph embeddings, we first apply GraphWave (Donnat et al., 2018) on the top-*K*-thresholded connectomes to produce a set of *d*-dimensional *node embeddings* for each of the *n* ROIs per connectome. GraphWave learns structural node embeddings, which individually capture the structural role of each node (ROI) within its local network neighborhood and in aggregate provide insights into the topological organization of the connectome graph. We then utilize principal components analysis (PCA) to summarize the set of *n d*-dimensional *node embeddings* into a single *graph embedding* vector by extracting the top 100 principal components. Aggregating these connectome graph embeddings across all time segments from all subjects results in a feature matrix of size m × 100.

**Table 1.** Symbols and abbreviations

| Symbol | Meaning | Value |
| --- | --- | --- |
| FC | Functional connectivity | - |
| dFC | Dynamic functional connectivity | - |
| WM | Working memory | - |
| ROI | Region of interest | - |
| GTD | Global temporal derivative | - |
| $N$ | Number of subjects | $N = 922$ |
| $n$ | Number of ROIs | $n = 268$ |
| $dt$ | Temporal derivative | - |
| $ts_i$ | Time series of ROI $i$ | - |
| $T$ | Length of time series | $T_{WM} = 405, T_{REST} = 1200$ |
| $t$ | Time point $t$ | - |
| $s, S$ | Time segment $s$ and total number of segments $S$, respectively | - |
| $C^{(s)}$ | Functional connectivity matrix for time segment $s$ | - |
| $ts_i(s)$ | Time series of ROI $i$ in time segment $s$ | - |
| $K$ | Number of edges retained in top-$K$ thresholding | $K = 10,000$ |
| $k$ | Number of clusters in $k$-means clustering | $k = (2 - 10)$ |
| $m$ | Total number of time segments/connectomes across all subjects in a single fMRI scanning session | $m_{WM1} = 8740, m_{WM2} = 9052$<br>$m_{REST1A} = 16,104, m_{REST1B} = 16,015$<br>$m_{REST2A} = 15,420, m_{REST2B} = 16,062$ |
| $d$ | Dimensionality of graph embedding | $d = 100$ |

We performed *k*-means clustering on the resultant group-level feature matrix, varying the number of clusters *k* in the range of 2–10. To determine the optimal number of clusters we utilized the elbow criterion of the cluster validity index, computed as the ratio of within-cluster distance to between-cluster distance (Allen et al., 2014). We mapped corresponding clusters across the session replicates to a single overall state based on shortest euclidean distances between the cluster centroid connectomes. Reproducibility of FC state clusters was tested across scanning sessions (two sessions for WM task, four sessions for resting state). Test-retest reliability was calculated across scanning sessions between centroids of corresponding states using the image intraclass correlation (I2C2) (Shou et al., 2013). We further characterize the resultant connectivity states with standard dFC features including average dwell time and state-to-state transition probabilities and go on to correlate these dFC features with neurophenotypes of interest.

### 2.3 Evaluation against ground truth

As described in Section 2.1.1, the WM task consists of four repeating task/rest blocks, where each block is structured as follows: 27.5s Task 1 (0-back), 27.5s Task 2 (2-back), 15s rest. This repeating task/rest block structure of the WM Task data serves as a natural ground truth for validation of our framework: if activation changes can truly be used as markers for connectivity changes, then we should be able to show that the discovered activation-informed change points align well with true onsets of WM task conditions. In fMRI data, signals are expected to be observed shortly after the stimulus, rather than directly aligned to the stimulus onset, due to lag in the hemodynamic response. Furthermore, the nature of block-design tasks results in sustained task-related activation changes rather than instantaneous spikes and subjects may require an additional 1-2s after the condition onset to fully enter the task state and experience the full effects of the task-induced activation response. Based on this, we defined a state change response window of 12 TR (8.6s) to account for the hemodynamic response time of 10 TR (7.2s) as well as an additional buffer of 2 TR (1.4s) for subjects to fully enter the task condition state. All peaks identified in the GTD series were labeled as either true positives or false positives based on whether they fell within the state change response window following a known task condition transition or not. Based on these labels, we calculate the overall precision and recall of our activation-informed change point detection, as well as the recall for transitions into each of the three task conditions (Task 1, Task 2, and Rest).

### 2.4 Comparison to Sliding Window

While the sliding window framework has been widely used to estimate dynamic FC states in resting fMRI where ground truth state changes cannot be known, it has not, to the best of our knowledge, been validated against a block-design task structure where the ground truth state changes are in fact known. To enable a direct comparison with the performance of our activation-informed segmentation method we applied the sliding window framework to the WM task data using the Group ICA of fMRI toolbox (GIFT) (https://trendscenter.org/software/gift/: Center for Translational Research in Neuroimaging and Data Science, Atlanta, Georgia) implementation, following the parameterization detailed in (Allen et al., 2014) as closely as possible. Specifically, we first performed group-level spatial independent component analysis (gICA) (Calhoun et al., 2001) to extract 50 independent components (ICs). IC time series then underwent a standard post-processing procedure to remove low-frequency trends associated with scanner drift, motion related variance and any other non-specific “spikes” or possible noise artifacts. Next, we utilized the dFNC function in the GIFT toolbox to perform the sliding window analysis. As in (Allen et al., 2014), we use a tapered window created by convolving a rectangle (size = 44 seconds/61 TR) with a Gaussian (σ = 3 TR) and sliding in steps of 1 TR, resulting in 344 total windows per WM fMRI session, and a total of 317,168 windows across all 922 subjects for each WM Session 1 and Session 2. Finally, the upper triangular of the windowed connectomes were used as feature vectors of length (50 × (49))/2 = 1225, and *k*-means clustering was applied to separate all windows into a set of *k* states. Due to the exceedingly long runtime of the *k*-means step on such a large dataset (317,168 × 1225), we manually set *k=* 3 for this sliding window analysis based on the three ground truth conditions in the block-design structure of this WM task.

We seek to quantify the accuracy of the resultant sliding window state clustering against the ground truth task conditions, in order to compare against the accuracy of our method proposed in this work. However, as we have noted in the introduction, the overlapping nature of the sliding window method makes interpretation of the resultant states difficult; in this case, most windows span both task and rest conditions so assigning a ground truth label to each window is not straightforward. Furthermore, there are no windows that only contain time points in the rest condition -in fact, in order to achieve this a window size of 15 seconds (21 TR) or less would be required, which is often regarded as too small for sliding window approaches. Considering this, we acknowledge that a “majority rules” labeling of windows would not be appropriate here, as all windows would be labeled as belonging to the task condition, so we devise two alternate labelings in order to be as fair as possible when evaluating the sliding window results. In the “full rest” labeling scheme we label only windows that contain the entirety of the rest block as belonging to the rest condition, and all other mixed windows as belonging to the task condition. In the “pure task” labeling scheme, we label only windows containing 90% or more task time points as belonging to the task condition, and all other windows containing >10% of resting time points as belonging to the rest condition (illustrated in Figure 4). Utilizing both of these ground truth labeling schemes enables us to test whether the sliding window paradigm is better suited to identifying pure task or majority rest windows from more mixed-condition windows. Lastly, due to the overlapping nature of the sliding windows identifying exact temporal change points between state conditions is not possible, therefore we cannot report precision and recall statistics for the sliding window framework.

## 3. Results

### 3.1. The GTD Method Accurately Identified Known Transitions During a Working Memory Task

Results of GTD-based peak discovery in WM task data are shown in Figure 3. The distribution of the discovered GTD peaks across all subjects showed a concentration of peaks immediately after a new condition onset (Figure 3B). In fMRI data, signals are expected to be observed shortly after the stimulus, rather than directly aligned to the stimulus onset, due to lag in the hemodynamic response. Using the true positive and false positive labels detailed earlier in Section 2.4, we found an average precision of 0.72 and average recall of 0.66 of all discovered change points against ground truth state transitions (Table 2). We found that Task 1 and Rest state onsets were more readily identifiable by our method than Task 2 onsets (Recall 0.67, 0.75, 0.57 respectively), indicating that transitions from task state to rest state and vice-versa elicit more significant changes in moment-to-moment activations than transitions from an easier 0-back WM task (Task 1) to a more difficult 2-back WM task (Task 2).

**Figure 3.**
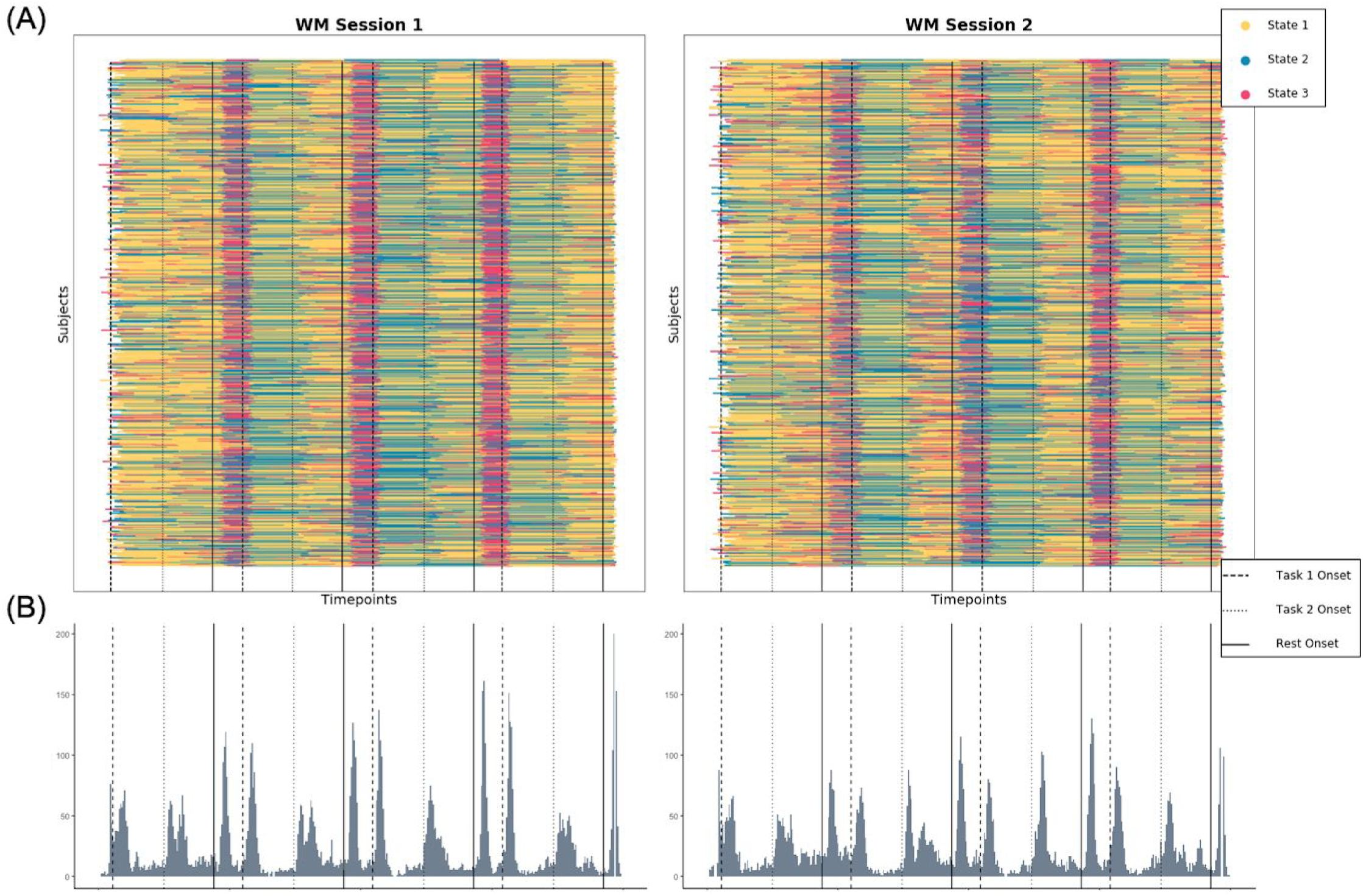
Results of the activation-informed segmentation for all subjects in structured WM task data. (A) Temporal alignment of our discovered segments colored by their corresponding state labels given by *k*-means clustering shows good alignment to known ground truth conditions (onsets marked by vertical lines: dashed for Task 1 onset, dotted for Task 2 onset, solid for Rest onset). (B) Histogram of discovered GTD peak locations show strong alignment to known condition onsets.

**Table 2.**
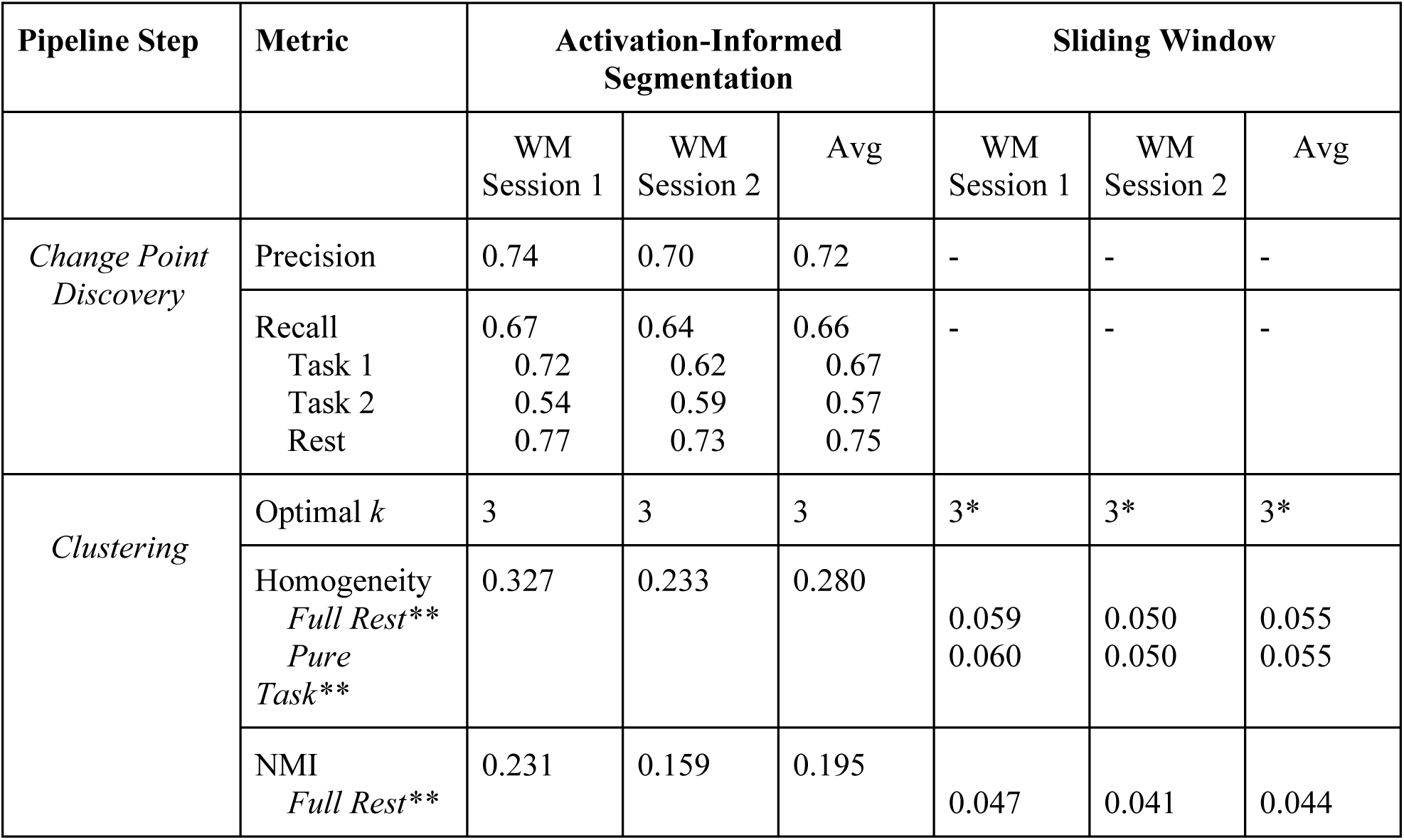
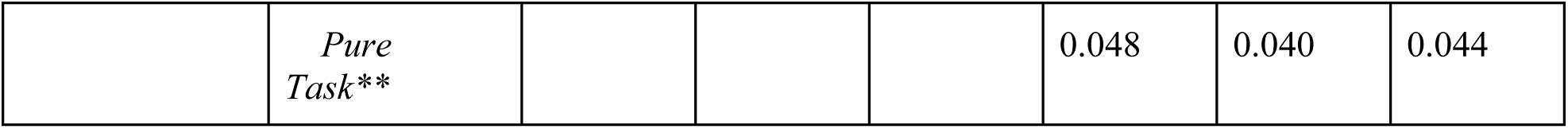
Performance of our activation-informed segmentation method and the standard sliding window method in recovering ground truth dynamic state changes in WM task data. The change point discovery step is unique to our framework and unable to be reported for the sliding window method. *Due to runtime constraints we manually set k = 3 in the GIFT sliding window approach rather than testing across a range of k values for the optimum **The overlapping nature of the sliding window necessitates choices to be made for ground truth labeling of windows, so we report performance against two ground truth labeling schemes specific to the sliding window (Section 2.4).

We found the optimal number of clusters *k =* 3 for both WM Session 1 and WM Session 2. Figure 3A illustrates the alignment of our segments, colored by their respective clusters, to the ground truth WM task conditions. Overall, we found good segregation between task and rest conditions, with improved accuracy in later block repetitions. As observed with the change point detection, the separation between Task 1 and Task 2 conditions is more difficult, owing both to the similarity in connectivity between the two working memory task conditions and to the lack of change point detection at Task 2 onset points resulting in segments that span the time frame of both Task 1 and Task 2. Homogeneity and normalized mutual information (NMI) metrics of our discovered clusters compared to the known ground truth are reported in Table 2. As our temporal segments may not directly align to the ground truth task blocks we derived ground truth labels for each discovered segment based on the corresponding task condition throughout the majority of the segment.

### 3.2. In the Working Memory Task, Activation-informed Segmentation Performance Was Superior to Sliding Window

We report the results of the GIFT toolbox sliding window pipeline for *k =* 3 states, which is most probable knowing the ground truth structure of the WM time series (Table 2). Though the sliding window approach does capture some repeating task versus rest signal (Figure 4), we found the GIFT sliding window approach had significantly decreased performance in segregating between known task and rest condition windows compared to our activation-informed segmentation approach, even when considering both the “full rest” and “pure task” ground truth labeling schemes (homogeneity = 0.055 vs. 0.280, respectively). Based on these results, we can conclude that our method more effectively and efficiently summarized the FC in each time segment, resulting in a 99.8% reduction in size of the final feature set passed to £-means compared to that of the sliding window approach (8740 × 100 vs. 317,168 × 1225 in WM Session 1). Furthermore, our method proved to be much more computationally efficient than the sliding window approach, and completing in < 2 hours for all subjects in a single WM session while the GIFT toolbox required > 24 hours to complete the requisite ICA and dFNC steps for the same data. Considering together the accuracy, data reduction and the runtime, we found our activation-informed segmentation method to be superior to the traditional sliding window paradigm in recovering dynamics in the context of a block-design ground truth.

**Figure 4.**
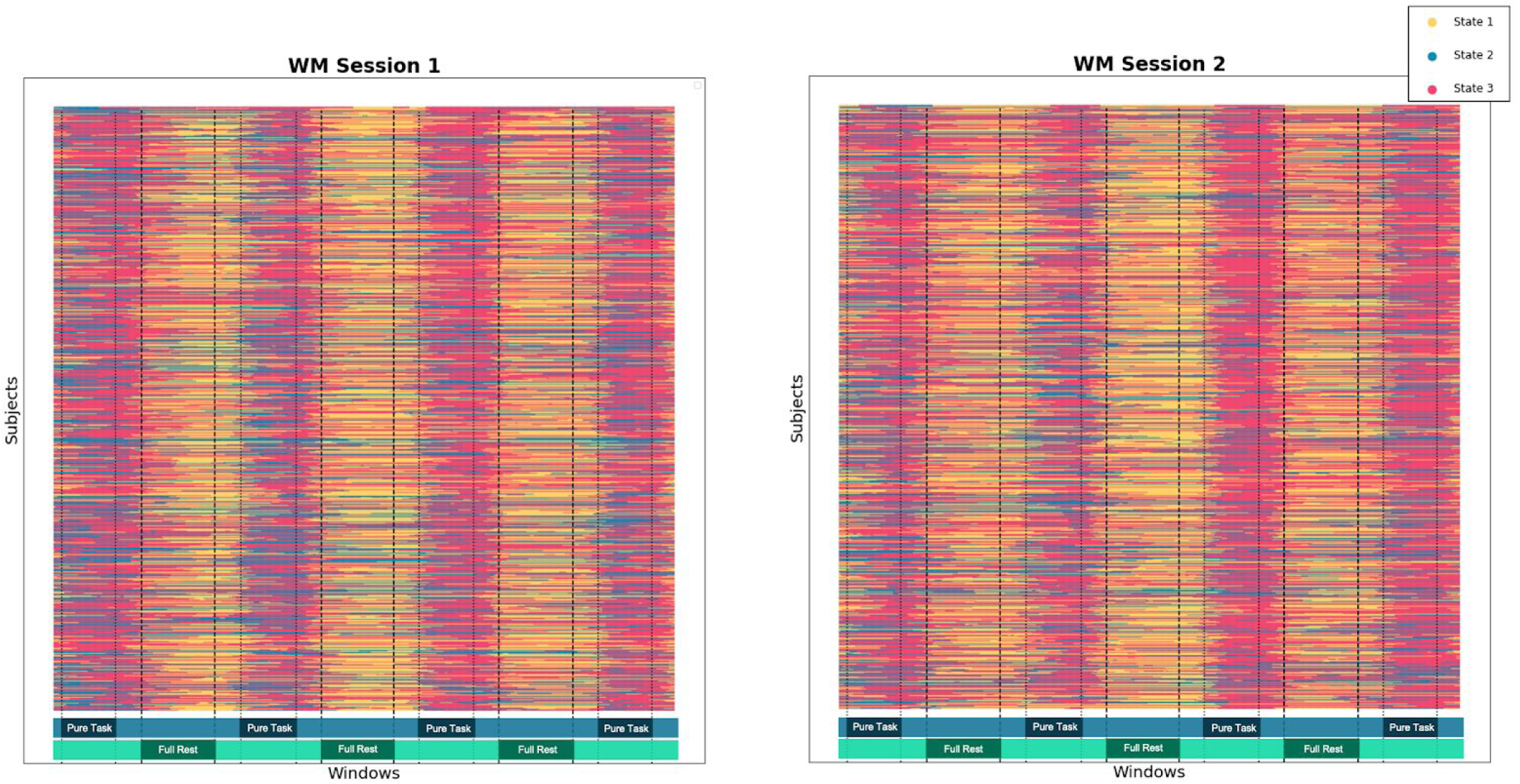
Results of GIFT toolbox-based sliding window framework for all subjects in structured WM task data. Both ground-truth labeling schemes (Section 2.4) are depicted with vertical lines (dashed for “full rest” and dotted for “pure task”) as well as colored bars along the lower horizontal axis.

### 3.3. The Activation-informed Segmentation Method Identified Five Connectivity States During Rest

We applied our activation-informed segmentation pipeline separately on four sessions of resting state fMRI data. Using the elbow criterion of the cluster validity index, we consistently found the optimal number of clusters *k =* 5 across the four sessions (Figure 5). Though our state clusters were derived using the graph embedding vectors as described above, we characterized the connectivity of each discovered cluster using the more interpretable top-*K* thresholded connectomes derived upstream in our pipeline for all segments in each cluster. We mapped corresponding clusters across the four session replicates to a single overall “dynamic state” based on shortest euclidean distances between the cluster centroid connectomes and found that each centroid was mapped only to one overall state by this criterion, indicating each state did indeed exhibit a unique connectivity signature.

**Figure 5:**
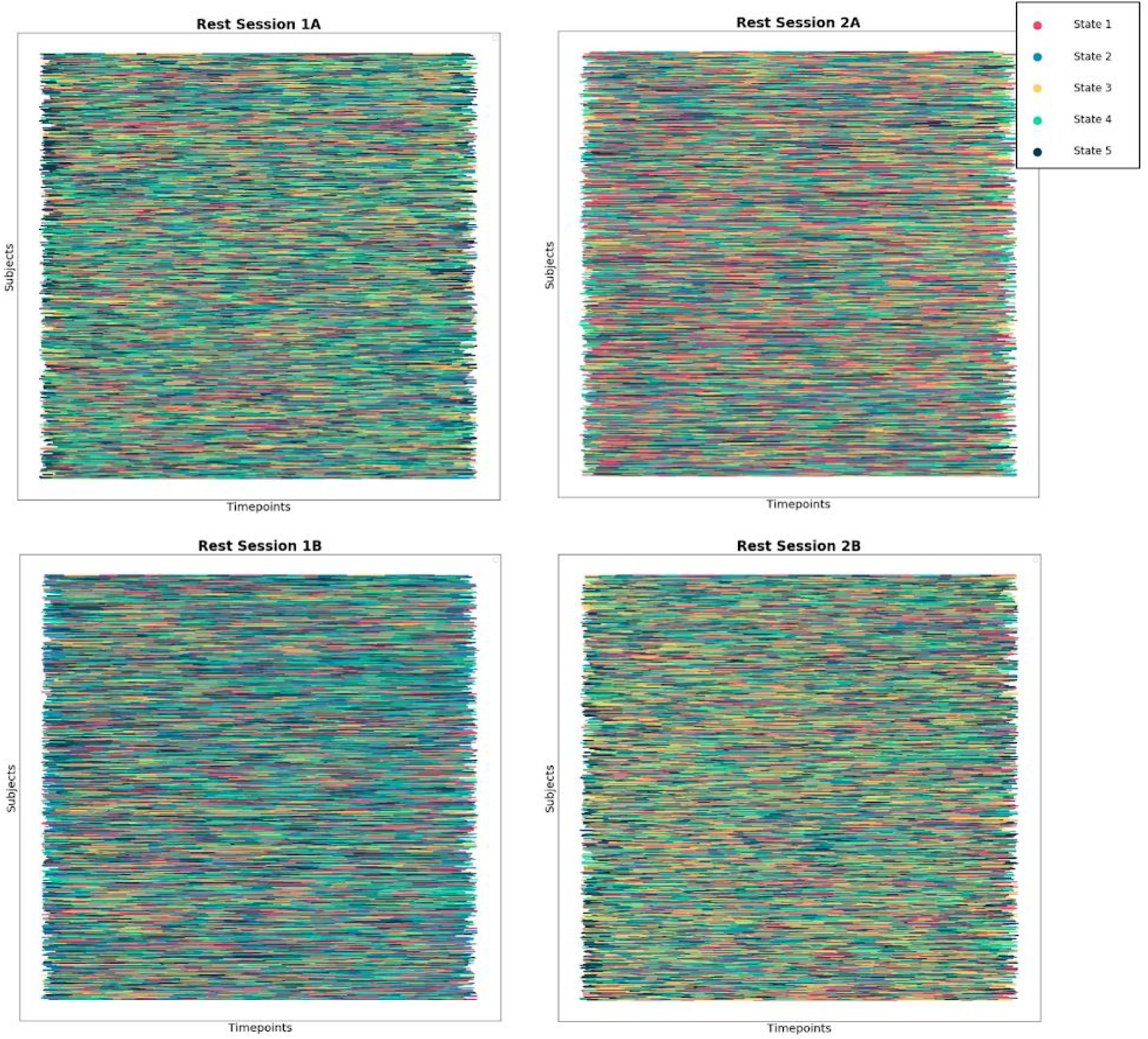
Temporal alignment of activation-informed segments and their corresponding state labels given by k-means in all four resting state fMRI sessions.

### 3.4. Connectivity States During Rest Exhibit Excellent Test-Retest Reliability

To assess the stability of these clusters we use the I2C2 metric, which was developed to assess the reliability of MRI images for a set of subjects across several image acquisition sessions. We consider each of the five dynamic states as “subjects” across each of the four resting state fMRI sessions for this calculation. We found very high replicability of our states across the four sessions (I2C2 = 0.96), suggesting that the dynamic states recovered by our method are indeed persistent across subjects and time, and may also be cognitively meaningful.

### 3.5. Activation Peaks Observed During Rest Closely Resemble Peaks Found When Transitioning In and Out of Cognitively Demanding Task States

We found that the magnitude of the GTD peaks that correspond to our discovered change points and define our dynamic states in rest are on the same order and mirror the distribution of the peaks found in the WM task setting (Kullback-Leibler divergence = 0.030) (Figure 6). This indicates that the changes in functional brain activity between dynamic states in rest are as strong as those observed when transitioning in and out of a cognitively demanding task state.

**Figure 6.**
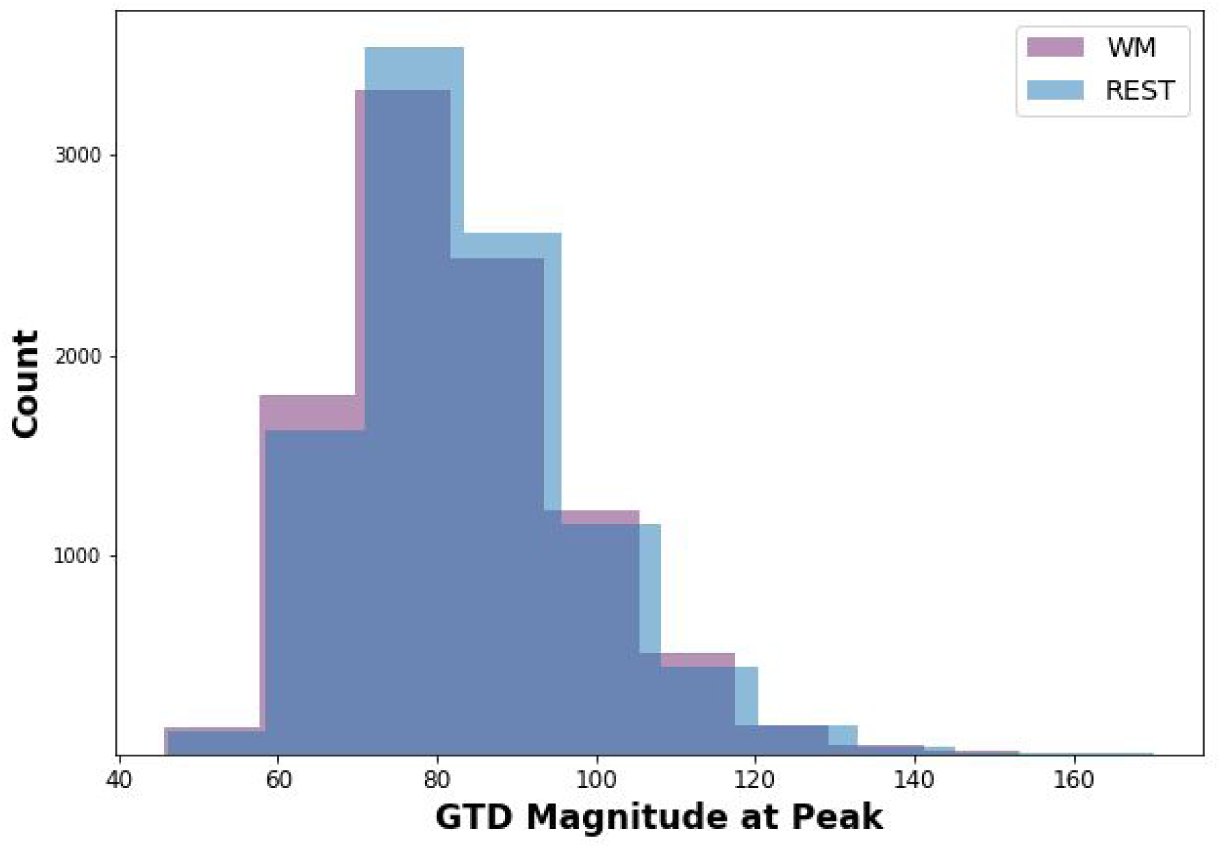
Histograms of GTD magnitudes at discovered peaks for 9700 change points in WM Session **1** and a size-matched random sample of change points in Rest Session 1A show similar distributions (Kullback-Leibler divergence = 0.030).

### 3.6. Connectivity States Involve Brain-Wide Connectivity Patterns and Prominently Involve Prefrontal/Sensory-Motor Coupling

We further characterized the overall connectivity signature of each resultant dynamic state by averaging the corresponding cluster centroids across the four sessions. This signature connectome for each of the five overall dynamic states is presented in Figure 7. Similar to results reported in (Nomi et al., 2017), states 1, 3, and 5 involve sensory/motor anti-correlation with the frontoparietal network. State 1 encompassed all sensory and motor networks, while state 3 had greater visual network specificity and state 5 had greater motor specificity. State 2 was characterized by anticorrelation between frontoparietal and medial frontal network, without sensory/motor involvement. State 4 exhibited none of the above motifs —just the within network connectivity that was common to all of the states.

**Figure 7.**
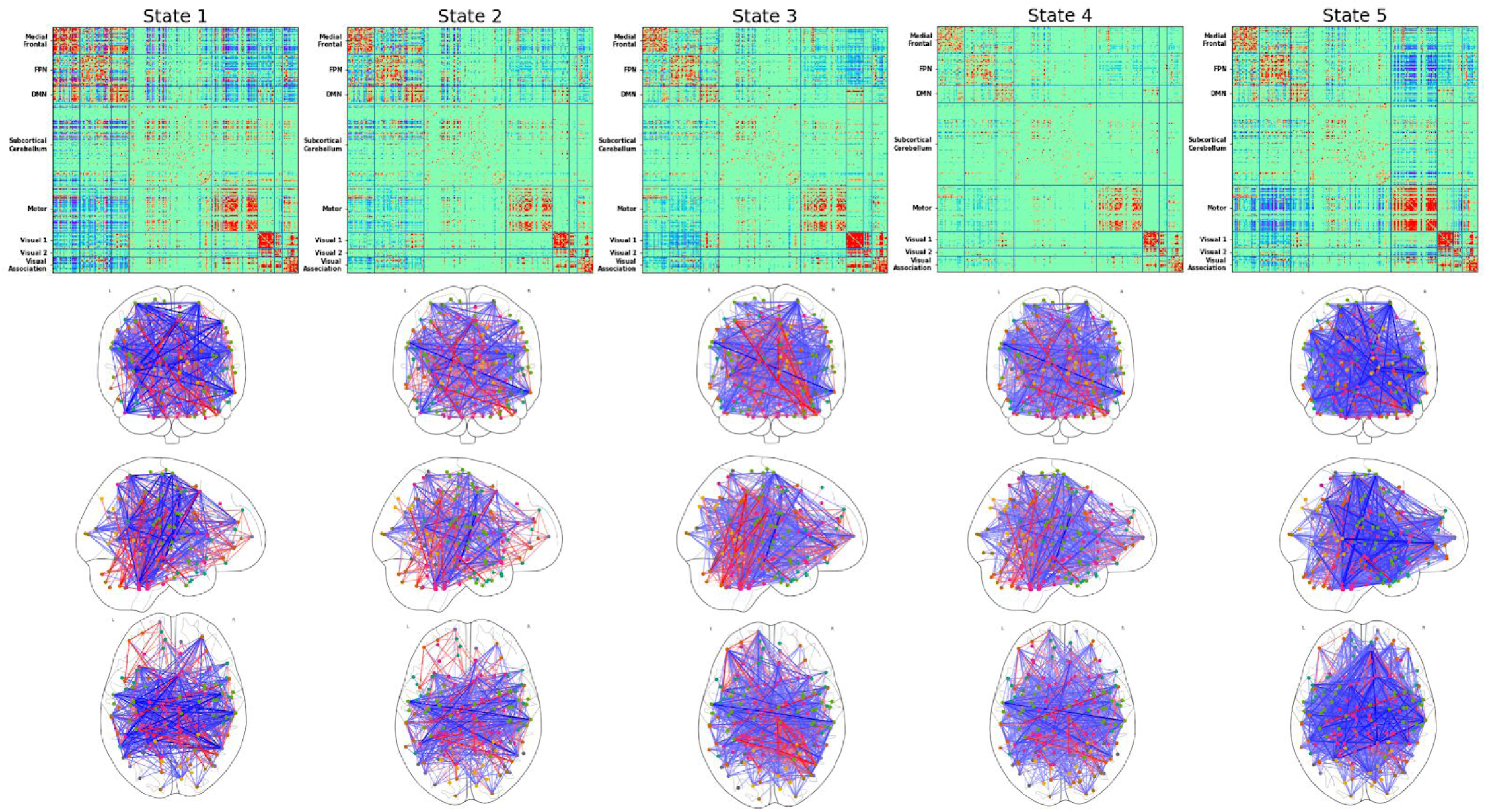
Connectivity signatures for each of our five discovered resting FC states. Connectivity signatures are defined by the centroid (i.e. average) of all connectomes belonging to each state cluster. Glass brain views show the top 0.5% of connections (360 edges) for each state.

### 3.7. Resting Connectivity States Exhibit Complex Patterns of Transitioning

In addition to summarizing each dynamic state by its unique connectivity patterns, we also extracted common dFC features including state-to-state transition probabilities, average dwell times per state, and number of occurrences of each state across the four resting sessions. We extracted these dFC features on a per-subject basis and then averaged them to capture the general patterns for all five dynamic states at the group level. The average state-to-state transition matrix, average dwell times, and average number of occurrences per state across all subjects are depicted in Figure 8. Overall, we found the highest probabilities of transitioning into state 4 from any of the other states. Interestingly, state 4 also exhibits the shortest dwell time of all five states, averaging a duration of 29.8 ±2.5 s, as well as the highest average number of occurrences. This coupled with the lower overall connectivity observed in state 4 suggests that this may represent a “buffer” state between the other dynamic states.

**Figure 8.**
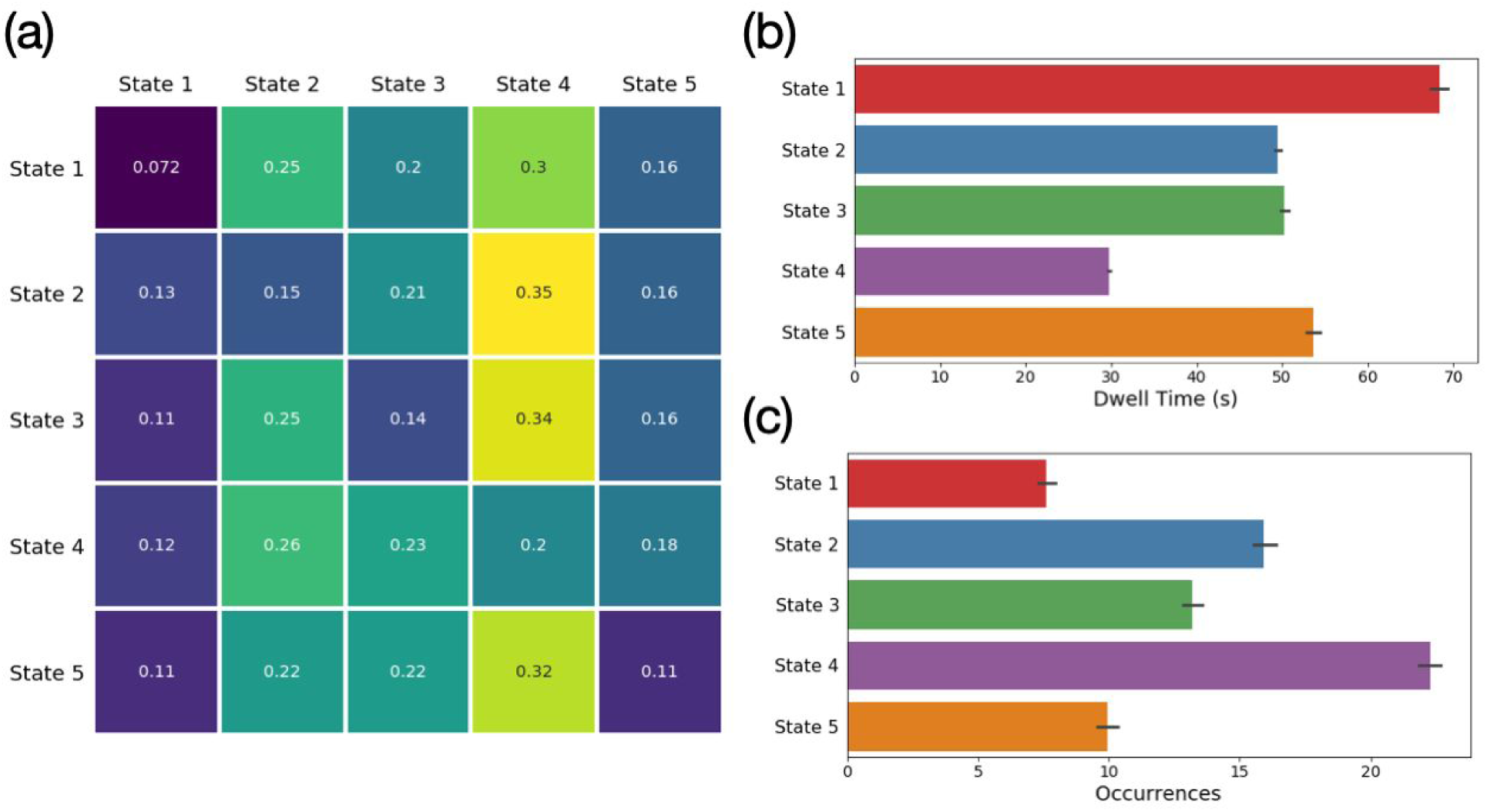
Average transition probabilities of moving from State A (along rows) to State B (along columns) (A), dwell times (B), and number of occurrences (C) across all subjects and resting state fMRI sessions.

### 3.8. Resting Connectivity States are Correlated with Behavioral Phenotypes Including Cognition, Personality, and Psychopathology

We performed correlation analysis between subject-specific dFC feature vectors, averaged across the four resting state sessions, and several neuro-relevant phenotypes. Specifically, we consider ten cognitive metrics: a general factor of intelligence (G; generated from a bifactor model as described in Sripada et al., 2020), processing speed (generated from factor modeling of three NIH Toolbox tasks as described in (Sripada et al., 2019)), the five facets of personality given by the Revised NEO Personality Inventory (openness to experience, conscientiousness, extraversion, agreeableness, and neuroticism), and the three dimensions of psychopathology given by the Adult Self Report Scale (Internalizing, Attention Problems, Externalizing). We report the ten strongest phenotype correlations in Table 3, along with corresponding uncorrected and false discovery rate (FDR) corrected p-values (Benjamini & Hochberg, 1995). We find several relatively strong correlations, three of which survive our stringent FDR corrected significance level (α = 0.05). Namely, we find significant relationships between the average dwell time in state 5 and conscientiousness (r = −0.1295; p = 0.0283), the number of occurrences of state 3 and processing speed (r = −0.1182; p = 0.0381), and the number of occurrences of state 4 and the general intelligence factor G (r = −0.1181; p = 0.0381).

**Table 3.** Ten strongest correlations of extracted dFC features to behavioral phenotypes of interest. Correlations that remain significant after FDR multiple test correction (α= 0.05) are denoted with (*).

| Feature | Phenotype | Correlation | Uncorrected p-value | FDR Corrected p-value |
| --- | --- | --- | --- | --- |
| Dwell Time State 5 | Conscientiousness | -0.1295 | 0.00008 | 0.0283* |
| Occurrence State 3 | Processing Speed | -0.1182 | 0.0003 | 0.0381* |
| Occurrence State 4 | G | -0.1181 | 0.0003 | 0.0381* |
| State 1 to State 5 Transition Probability | Agreeableness | 0.1074 | 0.0011 | 0.0956 |
| Dwell Time State 4 | G | -0.1038 | 0.0016 | 0.1122 |
| Probability of Remaining in State 1 | Conscientiousness | 0.1011 | 0.0021 | 0.1236 |
| State 3 to State 4 Transition Probability | G | -0.0937 | 0.0044 | 0.1948 |
| Occurrence of State 2 | Processing Speed | 0.0936 | 0.0045 | 0.1948 |
| State 1 to State 3 Transition Probability | Agreeableness | -0.0885 | 0.0072 | 0.2752 |
| Probability of Remaining in State 3 | Agreeableness | -0.0869 | 0.0083 | 0.2752 |

### 3.9. Resting Connectivity States are Unrelated to Head Motion

Head motion is a serious confound in studies of functional connectivity (Power et al., 2012; Satterthwaite et al., 2012; Van Dijk et al., 2012; Power et al., 2015). Moreover, it has recently been argued that head motion may in fact generate the time varying connectivity observed with sliding window methods (Laumann et al., 2017). We thus sought to determine whether the connectivity states we detected at rest with the GTD method were related to head motion. We found no significant correlation between the mean framewise displacement time series and the GTD series in all four resting state sessions (r = −0.0027; 95% CI = [−0.006, 0.0007]). We report all time-lagged cross-correlations for ± 10 TR in each of the four resting state sessions in Supplemental Table 2. This lack of correlation between framewise displacement and the GTD series suggests that there is no significant contribution of head motion to our discovered change points, and thereby our final dynamic states in rest. Taken together, these results strongly support the general existence of dynamicity in resting state and the reliability of the states discovered by our activation-informed framework.

## 4. Discussion

In this work, we introduce a new straightforward approach for assessing dynamic functional connectivity through informed time series segmentation. Our method, termed the activation-informed segmentation method, aims to derive FC states without the limitations of a predefined time scale for the dynamics or highly overlapped sliding windows. This framework is built upon the theory that changes in functional connectivity are mirrored by changes in functional activation. When applying the method to a working memory task where ground truth is known, we found that the method accurately found task boundaries, correctly recovered three connectivity states, and displayed a precision and recall profile that compared favorably to a leading sliding window approach. When applying the method to resting state data, we detected five connectivity states that displayed excellent test-retest reliability, exhibited complex transition dynamics, were correlated with multiple behavioral phenotypes, and were essentially unrelated to head motion. Our work expands the methodological toolkit for quantifying and characterizing time-varying connectivity and provides some of the strongest evidence to date for the existence of distinct dynamic states during rest.

We assessed the activation-informed segmentation method and sliding window approach head-to-head during a block-design working memory task to test whether these methods detect connectivity state changes where ground truth is known. Laumann et al. performed a test of the opposite issue: They examined a task with extended blocks where connectivity is assumed to be stable and found sliding window methods inappropriately found changing connectivity states where such changes are assumed to be absent (Laumann et al., 2017). In our test, the activation-informed segmentation method performed well. We observed an average precision of 0.72, meaning that 72% of activation changes detected by our algorithm corresponded to true changes in functional connectivity. Furthermore, the recall of true state transition points by our method averaged 0.66, and reached as high as 0.77 depending on the strength of the functional connectivity changes, indicating that a majority of known connectivity transitions are indeed marked by changes in global functional activation. In contrast, the GIFT sliding window method precludes the calculation of such precision and recall statistics due to the highly overlapping nature of the resultant windows. When considering the accuracy of the final state clustering, our method indeed performed ∼75% better than the sliding window method in separating blocks of true task from true rest. As far as we know, this is the first such test of the sliding window method in task data where ground truth is known. The fact that the sliding window has only fair accuracy in finding changes in connectivity state suggests there is room for improvement and reinforces our claim that further methods innovation in the study of time varying FC would be beneficial.

The activation-informed segmentation method found five states at rest and these states showed excellent test-retest reliability. These states appear to be broadly consistent with those reported in the previous literature in terms of number of states as well as connectivity patterns (Nomi et al., 2017). Furthermore, the mean dwell times are similar in duration. We also found these states are linked to a number of behavioral phenotypes -with the magnitudes of relationships similar to those reported in prior studies (Nomi et al., 2017). Taken together, these results suggest that there is some continuity in our results with the results from sliding window approaches. Nonetheless, some key differences remain. First, the states identified here have much higher test-retest reliability. Second, the method to identify them is simple, transparent, computed easily and efficiently, and appears not to be driven by artifactual causes (e.g., head motion).

A key assumption of our method is that activation changes can serve as a marker of changes in connectivity states. Several lines of evidence support this assumption. First, there is a substantial set of studies (discussed in the Introduction) that document connectivity patterns that arise during distinct task conditions. Importantly, these task conditions are antecedently known to produce distinct distributed activation profiles so that transitions into the relevant task conditions would produce activation shifts. Second, in the present study, we observed GTD peaks during the N-back working memory task when subjects shift task conditions, and we observed distinct connectivity states in the segments flanked by these GTD peaks. Third, if our main assumption were false, that is, if activation shifts *fail to* mark changes in connectivity states, then we should not have found large activation shifts during rest that are associated with distinct, highly test-retest reliable connectivity states. The fact that we did observe these results from rest provides further support that there is in fact a link between activation shifts and connectivity state changes. Finally, as we noted in the previous paragraph, the states identified have similarities along multiple dimensions with states identified through traditional sliding window methods. If our activation-informed segmentation approach can find connectivity states that are broadly similar to those found by sliding window approaches, this can only be explained if activation changes do indeed serve as a marker of connectivity changes.

In a somewhat unexpected finding, we observed GTD peaks during rest (corresponding to state change points) that were similar in magnitude to those seen during a working memory task. This finding is notable because the N-back working memory task is highly cognitively demanding and produces vigorous activations across a distributed “task-positive network” (Cabeza & Nyberg, 2000; Cole & Schneider, 2007; Mazoyer et al., 2001; Niendam et al., 2012). Rest, in contrast, is assumed to be a state of substantially reduced cognitive demands (McKiernan et al., 2003; Andrews-Hanna et al., 2010; Buckner et al., 2008). It is thus remarkable that we observed GTD peaks during the resting state on par with those that occur in response to transitions in and out of a cognitively demanding task. The fact that resting GTD peaks are so large provides additional support for our framework, which is based on the idea that easy-to-detect GTD peaks can be leveraged to identify hard-to-detect changes in connectivity states—large GTD peaks are particularly easy to detect. But critically, large GTD peaks during rest should be of independent interest to the field. That is, irrespective of their link to changes in connectivity states (which has been our focus in this study), the fact that there are regular and robust GTD peaks during rest is itself a phenomenon that needs follow up investigation and explanation.

There has been some skepticism in the field about the reality of time varying connectivity. A sizable portion of this debate centers on the sliding window methodology for demonstrating varying connectivity states (Laumann et al., 2017; Lindquist et al., 2014). It is claimed that this method generates artifacts, finds changes where none exist, etc. An important advance of the present study is that it demonstrates time-varying FC during rest without reliance on sliding window methods. Moreover, the associated connectivity states exhibit excellent test-retest reliability. Therefore, we believe that the present study offers the strongest evidence to date for the reality of time-varying connectivity at rest. More specifically, we suggest that the state transition points identified by our framework actually represent a lower bound of the “true” dynamic state changes in rest. This is because there is likely only an imperfect relationship between activation shifts and connectivity state changes: the former may be sufficient but not necessary for the latter. Thus, there may be at least some connectivity state changes that are not anteceded by prominent (and thus easy-to-detect) GTD peaks, and our method will fail to detect the presence of such connectivity states. Future work should seek to extend the change point detection scheme developed here to enable identification of these “connectivity-only” transitions. Such a method would require true moment-to-moment estimation of covarying activation patterns on a highly multivariate scale. It is possible these requirements can be fulfilled through the use of deep learning approaches, specifically recurrent neural network architectures, which are designed to learn complex, non-linear patterns in multivariate time series data (Li & Fan, 2018).

This study has several limitations. First, we rely on a key assumption that activation shifts serve as a marker for changes in connectivity states. We acknowledge that the relationship is likely imperfect and our method may underestimate the true number of states. The strength of our method, nonetheless, is simplicity and transparency, enabling the method to yield notably strong evidence for dynamic states at rest. Second, unlike sliding window methods that impose a uniform length on windowed connectivity matrices, the activation-informed segmentation method is sensitive to the duration of states. We mitigated this in multiple ways, including Fisher transformation and z-scoring of Pearson correlation-based connectivity matrices, as well as employing a top-*K* thresholding to control connectome density across both short and long segment lengths. Third, the meaning and importance of the dynamic states uncovered by the GTD method is unclear. We showed activation shifts are large (comparable to transitions in and out of a working memory task). We also presented initial data that connectivity states are linked to phenotypes of interest. But additional work is needed to establish what psychological processes are reflected in these dynamic states, and whether quantifying these transient states will yield significant theoretical and practical insights in psychology and neuroscience.

In sum, we introduce here a novel method for identifying dynamic states in fMRI without the use of potentially problematic sliding windows, validate the method in task data where ground truth is known, and demonstrate that the method finds strong evidence—likely the strongest evidence to date—for the presence of dynamic states at rest.

## Acknowledgments

This work was supported by funding from the University of Michigan Precision Health Investigator Award. MD was supported by the National Science Foundation Graduate Research Fellowship Program grant DGE-1256260. The authors thank Parmida Davarmanesh, Aman Taxali, Saige Rutherford and Mike Angstadt for their support and valuable feedback throughout this work.

---

**Supplemental Table 1.** Clustering performance of traditional FC summarization methods in our activation-informed segments.

| Method | Optimal $k$ | | Homogeneity | | NMI | |
| --- | --- | --- | --- | --- | --- | --- |
|  | *WM*<br>*Session 1* | *WM*<br>*Session 2* | *WM*<br>*Session 1* | *WM*<br>*Session 2* | *WM*<br>*Session 1* | *WM*<br>*Session 2* |
| Connectome Vectorization | 3 | 3 | 0.036 | 0.037 | 0.025 | 0.025 |
| PCA Decomposition | 3 | 3 | 0.033 | 0.035 | 0.024 | 0.024 |

**Supplemental Table 2.** Mean correlation and 95% confidence interval (CI) between GTD and framewise displacement time series at lags ranging from (−10,10) TR for four sessions of resting state fMRI.

| Session | Lag | Mean Correlation | 95% CI Lower Bound | 95% CI Upper Bound |
| --- | --- | --- | --- | --- |
| REST1B | -10 | -0.0063 | -0.0100 | -0.0025 |
| REST1B | -9 | -0.0089 | -0.0127 | -0.0052 |
| REST1B | -8 | -0.0126 | -0.0164 | -0.0088 |
| REST1B | -7 | -0.0174 | -0.0213 | -0.0135 |
| REST1B | -6 | -0.0221 | -0.0262 | -0.0181 |
| REST1B | -5 | -0.0259 | -0.0301 | -0.0217 |
| REST1B | -4 | -0.0272 | -0.0315 | -0.0229 |
| REST1B | -3 | -0.0260 | -0.0304 | -0.0216 |
| REST1B | -2 | -0.0228 | -0.0273 | -0.0184 |
| REST1B | -1 | -0.0189 | -0.0233 | -0.0145 |
| REST1B | 0 | -0.0152 | -0.0195 | -0.0109 |
| REST1B | 1 | -0.0133 | -0.0176 | -0.0090 |
| REST1B | 2 | -0.0135 | -0.0178 | -0.0093 |
| REST1B | 3 | -0.0149 | -0.0192 | -0.0107 |
| REST1B | 4 | -0.0160 | -0.0202 | -0.0118 |
| REST1B | 5 | -0.0152 | -0.0195 | -0.0110 |
| REST1B | 6 | -0.0122 | -0.0164 | -0.0080 |
| REST1B | 7 | -0.0071 | -0.0112 | -0.0029 |
| REST1B | 8 | -0.0009 | -0.0049 | 0.0031 |
| REST1B | 9 | 0.0050 | 0.0010 | 0.0090 |
| REST1B | 10 | 0.0095 | 0.0054 | 0.0135 |
| REST1A | -10 | 0.0039 | 0.0008 | 0.0071 |
| REST1A | -9 | 0.0035 | 0.0004 | 0.0067 |
| REST1A | -8 | 0.0028 | -0.0003 | 0.0060 |
| REST1A | -7 | 0.0021 | -0.0010 | 0.0052 |
| REST1A | -6 | 0.0016 | -0.0015 | 0.0047 |
| REST1A | -5 | 0.0014 | -0.0017 | 0.0045 |
| REST1A | -4 | 0.0018 | -0.0013 | 0.0050 |
| REST1A | -3 | 0.0022 | -0.0009 | 0.0053 |
| REST1A | -2 | 0.0021 | -0.0010 | 0.0052 |
| REST1A | -1 | 0.0018 | -0.0013 | 0.0049 |
| REST1A | 0 | 0.0016 | -0.0015 | 0.0048 |
| REST1A | 1 | 0.0014 | -0.0018 | 0.0045 |
| REST1A | 2 | 0.0010 | -0.0021 | 0.0041 |
| REST1A | 3 | 0.0008 | -0.0023 | 0.0038 |
| REST1A | 4 | 0.0008 | -0.0022 | 0.0038 |
| REST1A | 5 | 0.0013 | -0.0017 | 0.0043 |
| REST1A | 6 | 0.0020 | -0.0010 | 0.0049 |
| REST1A | 7 | 0.0027 | -0.0002 | 0.0057 |
| REST1A | 8 | 0.0035 | 0.0006 | 0.0064 |
| REST1A | 9 | 0.0040 | 0.0011 | 0.0069 |
| REST1A | 10 | 0.0042 | 0.0013 | 0.0071 |
| REST2B | -10 | 0.0029 | -0.0001 | 0.0060 |
| REST2B | -9 | 0.0028 | -0.0002 | 0.0059 |
| REST2B | -8 | 0.0028 | -0.0003 | 0.0059 |
| REST2B | -7 | 0.0027 | -0.0004 | 0.0058 |
| REST2B | -6 | 0.0024 | -0.0006 | 0.0055 |
| REST2B | -5 | 0.0020 | -0.0010 | 0.0051 |
| REST2B | -4 | 0.0017 | -0.0013 | 0.0047 |
| REST2B | -3 | 0.0013 | -0.0018 | 0.0043 |
| REST2B | -2 | 0.0005 | -0.0026 | 0.0036 |
| REST2B | -1 | 0.0000 | -0.0031 | 0.0031 |
| REST2B | 0 | -0.0001 | -0.0031 | 0.0030 |
| REST2B | 1 | 0.0001 | -0.0030 | 0.0031 |
| REST2B | 2 | 0.0001 | -0.0030 | 0.0032 |
| REST2B | 3 | 0.0000 | -0.0030 | 0.0030 |
| REST2B | 4 | -0.0001 | -0.0031 | 0.0029 |
| REST2B | 5 | -0.0001 | -0.0030 | 0.0029 |
| REST2B | 6 | 0.0000 | -0.0030 | 0.0029 |
| REST2B | 7 | 0.0002 | -0.0028 | 0.0031 |
| REST2B | 8 | 0.0005 | -0.0025 | 0.0035 |
| REST2B | 9 | 0.0007 | -0.0022 | 0.0037 |
| REST2B | 10 | 0.0007 | -0.0022 | 0.0037 |
| REST2A | -10 | 0.0009 | -0.0021 | 0.0038 |
| REST2A | -9 | 0.0009 | -0.0021 | 0.0039 |
| REST2A | -8 | 0.0013 | -0.0017 | 0.0042 |
| REST2A | -7 | 0.0015 | -0.0014 | 0.0044 |
| REST2A | -6 | 0.0018 | -0.0011 | 0.0047 |
| REST2A | -5 | 0.0019 | -0.0010 | 0.0048 |
| REST2A | -4 | 0.0020 | -0.0009 | 0.0049 |
| REST2A | -3 | 0.0021 | -0.0009 | 0.0050 |
| REST2A | -2 | 0.0020 | -0.0010 | 0.0050 |
| REST2A | -1 | 0.0022 | -0.0008 | 0.0053 |
| REST2A | 0 | 0.0029 | -0.0002 | 0.0059 |
| REST2A | 1 | 0.0035 | 0.0004 | 0.0065 |
| REST2A | 2 | 0.0038 | 0.0007 | 0.0068 |
| REST2A | 3 | 0.0040 | 0.0010 | 0.0070 |
| REST2A | 4 | 0.0041 | 0.0011 | 0.0071 |
| REST2A | 5 | 0.0043 | 0.0014 | 0.0073 |
| REST2A | 6 | 0.0046 | 0.0016 | 0.0075 |
| REST2A | 7 | 0.0048 | 0.0019 | 0.0078 |
| REST2A | 8 | 0.0050 | 0.0021 | 0.0080 |
| REST2A | 9 | 0.0050 | 0.0020 | 0.0080 |
| REST2A | 10 | 0.0048 | 0.0018 | 0.0078 |

